# Evaluation of the morphological and biological functions of vascularized microphysiological systems with supervised machine learning

**DOI:** 10.1101/2023.01.12.523755

**Authors:** James J. Tronolone, Tanmay Mathur, Christopher P. Chaftari, Abhishek Jain

## Abstract

Vascularized microphysiological systems and organoids are contemporary preclinical experimental platforms representing human tissue or organ function in health and disease. While vascularization is emerging as a necessary physiological organ-level feature required in most such systems, there is no standard tool or morphological metric to measure the performance or biological function of vascularized networks within these models. Further, the commonly reported morphological metrics may not correlate to the network’s biological function – oxygen transport. Here, a large library of vascular network images was analyzed by the measure of each sample’s morphology and oxygen transport potential. The oxygen transport quantification is computationally expensive and user-dependent, so machine learning techniques were examined to generate regression models relating morphology to function. Principal component and factor analyses were applied to reduce dimensionality of the multivariate dataset, followed by multiple linear regression and tree-based regression analyses. These examinations reveal that while several morphological data relate poorly to the biological function, some machine learning models possess a relatively improved, but still moderate predictive potential. Overall, random forest regression model correlates to the biological function of vascular networks with relatively higher accuracy than other regression models.

## 1. Introduction

Organ-chips, also known as microphysiological systems (MPS), are an *in vitro* research tool with increasing applications in pharmaceutics testing/evaluation^1-3^, tissue engineering and disease modeling^4-6^, and medical regulation^7,8^. Recent success of MPS has been reported in evaluating drug candidates and therapies for rapidly arising diseases^9^, predicting outcomes of patient-specific therapies^10-12^, and even their rapidly increasing complexity with increasing similarity to human tissues and organs^13-16^. An emerging direction within the MPS development is the vascularization of these systems^17,18^, where perfusable vasculature anywhere from small capillary beds to large veins and arteries are modeled with human tissues, blood, and biophysical stimuli^19-22^. Vascularized MPS (termed as vMPS) may include the self-assembly of small capillary networks around other cell types, such as cancer organoids, pancreatic islets, lung epithelia, and brain/neural cells^18^.

Typically, vMPS are engineered by applying a combination of physical and chemical stimulants of network formation^23,24^, for example, endothelial cell source and density, proangiogenic/supporting cell source and density, extracellular matrix, biochemical cues, and biophysical stimuli^25^. Despite the diversity of ways vMPS may be constructed, the ultimate outcome of these systems is to achieve a vascular bed formation in a tissue microenvironment that mimics an *in vivo* state. However, we observed that published reports use different evaluation criteria of the network quality and function based on a range of quantification tools, and there is no consistency. These methods mostly quantify the morphology of the microvascular networks, but the morphology’s relationship to the network’s actual function, transport of oxygen to deoxygenated tissues, remains unclear. This has prevented a standardization of the field that could help its translation to large pharmaceutical, tissue engineering, or clinical studies^26^.

Our work here assesses the ability of a vascular network to transport oxygen and its correlation with most common morphological metrics that have been reported in the past. Using a large library of images with diverse formations of vMPS, we measured the morphological metrics according to previous *in vitro* vasculature evaluation methods, as well as computed the biological function of the networks through measures of Normalized Oxygen Flux (NOF), Normalized Vascular Potential (NVP), and Normalized Oxygen Delivery (NOD). Since an exponentially increasing number of advanced statistical and machine learning tools are now readily available to the casual programmer, we have applied several of the most relevant techniques – multiple linear regression, tree-based regression including random forest – to discover relationships between biological and morphological metrics of vMPS. Our analyses identify several redundant and multicollinear metrics, data handling practices, as well as algorithms consisting a set of few most relevant independent metrics that may be adopted for quantitative analyses of vMPS.

## 2. Methods

### Microfluidic Device Fabrication

Microfluidic devices were manufactured according to traditional photo- and soft-lithography practices. Detailed methods are listed in the supplementary material. Briefly, photomasks containing multiple iterations of the microfluidic device were designed using Solidworks 2019 (Dassault Systems) and printed onto mylar films (CAD-Art Services; Bandon, Oregon, USA). There were used for exposing UV light to 250 μm layers of SU-8 (Kayaku Advanced Materials; Westborough, MA 01481) on 6-inch silicon wafers (University Wafer; South Boston, MA 02127, USA). Exposed wafers were developed (Kayaku Advanced Materials) and subsequently silanized with trichloro(1H,1H,2H,2H-perfluorooctyl)silane (PFCOTS) (448931-10G, Sigma Aldrich) overnight.

Polydimethylsiloxane (PDMS) (Dow Corning) was prepared in a 10:1 base:curing agent ratio, poured over the wafer, degassed, and baked at 70 °C for 2 hr. Cured PDMS was peeled from the master mold and port holes were punched. Devices and glass slides were placed in a 100 Watts plasma cleaner (Thierry Zepto, Diener Electronics; Ebhausen 72224, Germany), treated, bonded together irreversibly, and then left at 70 °C overnight to render the PDMS hydrophobic. Before cell culture, devices were left under UV for 30 min for sterilization.

### Cell Culture

Human umbilical vein endothelial cells (HUVECs) (C2519A, Lonza; Houston 77047, TX) cultured in endothelial growth medium-2 (EGM-2) (C-22111, Promocell; Heidelberg 69126, Germany), human microvascular endothelial cells (HMVECs) (CC-2527, Lonza) cultured in endothelial growth medium-2 MV (EGM-2 MV) (CC-22121, Promocell), and normal human lung fibroblasts (NHLFs) (CC-2512, Lonza) cultured in fibroblast growth medium (FGM) (CC-3132, Lonza) were used from passages 4-6. After expansion, cells were trypsinized, pelleted, and resuspended in media for counting and viability assessment. Cell suspensions were adjusted to desired concentrations and mixed with thrombin (605157-1KU, Sigma Aldrich) at a final concentration of 5 U/mL. Fibrinogen (F8630-1G, Sigma Aldrich) was dissolved in 1X PBS, filtered, and supplemented with 0.3 mg/mL collagen I (354236, Corning) and 0.2 U/mL aprotinin (A3248-10MG, Sigma Aldrich). 2X fibrinogen and 2X thrombin/cell solutions were mixed 1:1 and injected into hydrogel compartments. Fresh EGM-2 was injected into the fluidic channels, followed by daily media changes for the duration of the 96 hr culture.

### Immunostaining

Devices were fixed with 4% paraformaldehyde (AAJ19943K2, Fisher Scientific) for 15 minutes at room temperature, washed, and permeabilized with 0.1 vol% TritonX-100 (T8787-100ML, Sigma Aldrich) in 3% bovine serum albumin (BSA) (BP9706100, Fisher Scientific) for 15 minutes at room temperature. Devices were washed, then stained using AlexaFluor-488-conjugated CD-31 antibody (303110, Biolegend; San Diego, CA 92121, USA) diluted 1:100 in 3% BSA overnight at 4 °C. Devices were washed and stained with rhodamine phalloidin (R415, Thermo Fisher) for 1 hour at room temperature, washed, and counterstained with Hoescht (H2570, Thermo Fisher) for 10 minutes at room temperature.

The vascularization region of each sample was imaged using Z-stack and tiling acquisition with a Zeiss Axio Observer Z1 Inverted Microscope (Zeiss; Thornwood, New York 10594, USA). Stitched images were orthogonally projected in the XY direction and cropped to and area of 1×1 mm.

### Image Processing and Vessel Morphology Quantification

Sectioned images of each sample’s vascularization region were processed using REAVER^27^. All images were segmented using user-defined parameters (filter size = 64; minimum connected component area = 800; wire dilation threshold = 0; vessel thickness threshold = 1; gray threshold = 0.045). Images were quantified and measurements for all samples were compiled in GraphPad Prism 9 (GraphPad Software, Inc.; San Diego, CA 92180, USA) datasheets.

### Numerical Analysis

Detailed coding methods are discussed in the supplementary material. Briefly, vMPS images were imported to MATLAB, converted to triangular meshes for the hydrogel and vasculature regions, and composited. AngioMT detected the vessel edges and applied the oxygen boundary conditions of a cell culture incubator. AngioMT added an oxygen permeability flux at the edges and solved a steady-state, two-dimensional transport of oxygen for each node within the vessel, and then within the tissue. The nodal oxygen concentrations were summarized as area averages and normalized to the input concentration.

### Data Analysis and Machine Learning

All data analyses were carried out using Python 3.8.8 via Anaconda3, and all code was written in JupyterLab 3.0.14 web browser. REAVER datasets were modified with an additional column for NOF, NVP, and NOD measurements corresponding to each sample and used as input for all analyses.

Detailed coding methods are discussed in the supplementary material. Briefly EDA was begun by generating histograms, correlation heatmaps, and scatter plots. VIF was calculated to quantify the degree of multicollinearity by

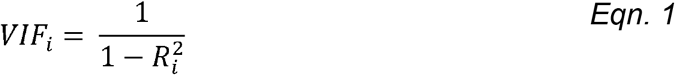

where *i* represents the index of an independent variable, and *R*^*2*^ is the coefficient of determination.

Feature engineering began with PCA, first by scaling morphological data. For visualization’s sake, the new PCs were segregated based on their input NOD value: Great – 1.0-0.75; Good – 0.50-0.75; OK – 0.25-0.50; Bad – 0.0-0.25 and plotted against PC1-2. Explained variances, eigenvalues, and loadings were obtained from the PCA object, from which scree plots were made. Heatmaps of PCs vs. metrics were generated as previously described.

Input data for FA was scaled similar to the PCA methods. Bartlett Sphericity and KMO tests were generated to test for analysis appropriateness. Unrotated FA was first carried out using 6 factors, followed by Varimax and Promax rotations. Heatmaps of all Factors 1-2 vs. inputs were generated as previously described.

Data for machine learning was split into training and testing sets using with an 80-20% split. Linear regression was used to fit the training dataset to an ML model. The testing set was predicted with the fit model and evaluated with traditional accuracy metrics (R^2^, MAE, MSE, RMSE). Lines of best fit were plotted over scatter plots of the expected vs. predicted values in order to show the prediction variances. RFECV was used to eliminate least-impacting inputs until the R^2^ value decreased.

Decision tree regression was carried out followed with pruning via cost complexity analysis after determining an optimal effective alpha value. Random forest models were generated using 100 estimators. Feature importances were represented as decreases in accuracy following morphological metric perturbation.

All plots were exported as SVG files and compiled into publication-ready figures using Adobe Illustrator (Adobe, Inc.; San Jose, CA 95110, USA).

## 3. Results

### Generation of microvascular networks for quantitative standardization and evaluation

To acquire a broad range and diversity of microvascular networks, we adopted a relatively standard microfluidic device from previous literature that allowed the compartmentalization of alternating hydrogel and fluid channels (**Figure 1A-D**)^28-31^. This compartmentalization facilitated the diffusion of paracrine signals from co-cultured cells, or allowed exogenous supplementation of growth factors in cell culture medium, all of which yielded perfusable vascular networks within 96 hours (**Figure 1D**). In most assemblies, proangiogenic fibroblasts (LFs) were situated in outermost hydrogel compartments, but other assemblies in the dataset included mono-cultures of ECs in central compartments and acellular fibrin gels in outermost channels, the same supplemented with growth factors sourced through media channels, and even fibroblasts co-cultured within the central channel with ECs to facilitate LF-EC cell-cell contacts. In all cases, microvascular networks self-assembled and Z-stack images were compiled, cropped, and prepared for downstream, quantitative analysis.

**Figure 1.**
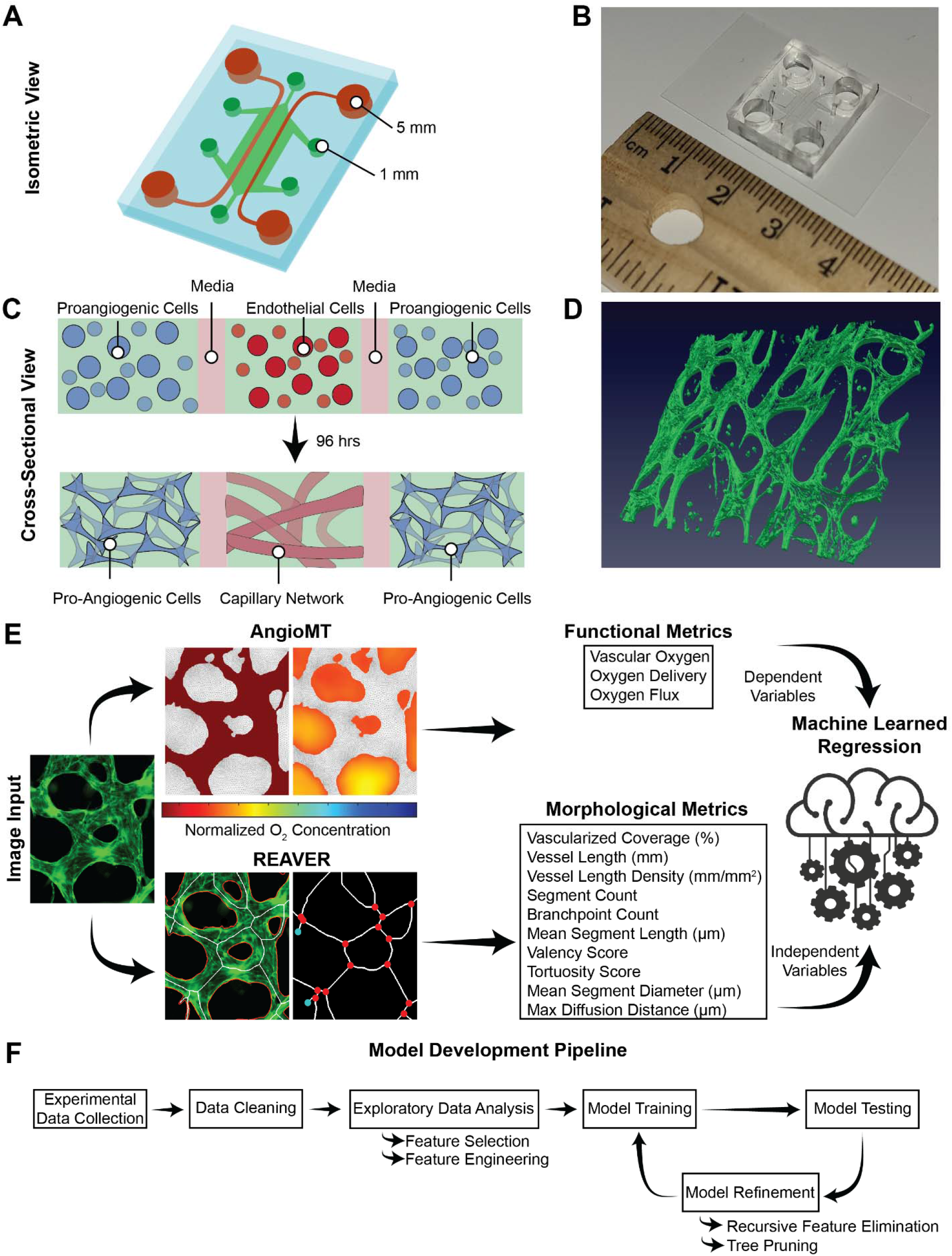
Machine learning problem definition and development pipeline. **A**) schematic showing the parallel situation of five microchannels terminated in 1- or 5-mm port holes for hydrogel and fluid channels, respectively. **B**) fabricated device. **C**) cell suspensions are encapsulated in hydrogels and injected into the vMPS. After 96 hours, endothelial cells form tubular networks similar to an in vivo capillary bed. **D**) 3D reconstruction of a vMPS. **E**) input images are processed with AngioMT and REAVER, which give biological/functional and morphological quantitations, respectively. **F**) The model development pipeline including data exploration and analysis via feature selection and engineering, as well as model refinement through cost-complexity analyses that eliminate insignificant input variables.

Traditional vascular network evaluation methods include the segmentation, skeletonization, and quantification of individual morphological metrics such as vessel length (mm), diameter (μm), and coverage (%)^26^, described in **Table S1**. While these metrics quantitate morphological features of the network, it is unclear if they relate to the biological function of these networks. In a recent work, we created an *in silico* platform, named as AngioMT, that predicts oxygen transport phenomena within these networks (Mathur, et al. *unpublished*). Since a vascular network’s biological performance may be dependent upon the end-to-end movement of oxygen across a network, leakiness or permeability of capillary networks, and actual oxygen delivery into surrounding parenchyma, we derived metrics that quantitated these phenomena and measured them in our AngioMT software. We derived a Normalized Vascular Potential (NVP) as a measure of permeability and the amount of oxygen available to be delivered by tissues; a Normalized Oxygen Delivery (NOD) as a measure of the oxygen delivery into hydrogel regions, calculated as the area average of oxygen over all regions representing deoxygenated tissue, and a Normalized Oxygen Flux (NOF) as a measure of the average oxygen diffusion rate from vessels to tissues describing volumetric transport of oxygen in the tissue regions (**Table S1**).

While these biological descriptors are directly related to the oxygen tension of vascular networks, the computational cost, time, lack of automated analyses and the requirement of prior user expertise limits AngioMT’s utilization. For example, calculating these biological parameters may require significant image processing over several hours and the process may be user-dependent and not entirely objective. Therefore, due to AngioMT’s high computational cost, and REAVER’s unexplained physiological relevance, we aimed to develop a machine learned regression predicting a vMPS’s function from its morphology. We were inspired to develop a machine learned regression model from which low complexity morphological quantifications could be used to predict the higher complexity functional quantifications (**Figure 1E**) using established algorithms (**Figure 1F**) for rapid, user-independent and automated outcomes.

### Metrics of vascular network function: Statistical tests of independence and multicollinearity

Since all the morphological and biological metrics (**Table S1**) describe different aspects of the vascular network, the extent to which they are specific, independent, and physiologically relevant is unknown. Further, inclusion of unnecessarily high dimensional data may require unnecessary computational time and redundancies in the machine learning analysis. Since several variables describe the vascular networks (**Table S1**), we began an exploratory data analysis (EDA) to select an optimal number of input features with the goal of reducing the high dimensionality of our dataset and to reduce the model training time. In our vascular networks (vMPS), we first analyzed the variable distribution and interquartile range between their minima and maxima (**Table S2**), as well as each variable’s proportion of outliers (**Table S3**), finding that mean tortuosity and mean valency skewed towards a single value. This was confirmed upon an examination of the distributions of density of each morphological variable (**Figure 2A**). Further, because mean tortuosity and mean valency required other morphological values for computation (**Table S1**), we opted to exclude these two metrics from the machine learning analysis. Additionally, we found that max diffusion distance had a high proportion of outliers (**Table S3**), and thus, may also be excluded. We also found that vessel length density was similar in scale to the vessel length (**Figure 2B**), and therefore, using both as input variables is redundant in our downstream regression analyses. In summary, our analyses showed that vessel coverage, vessel length, branchpoint count, segment count, mean segment length, and mean segment diameter are statistically suitable as input variables for machine learning algorithm development and regression analysis.

**Figure 2.**
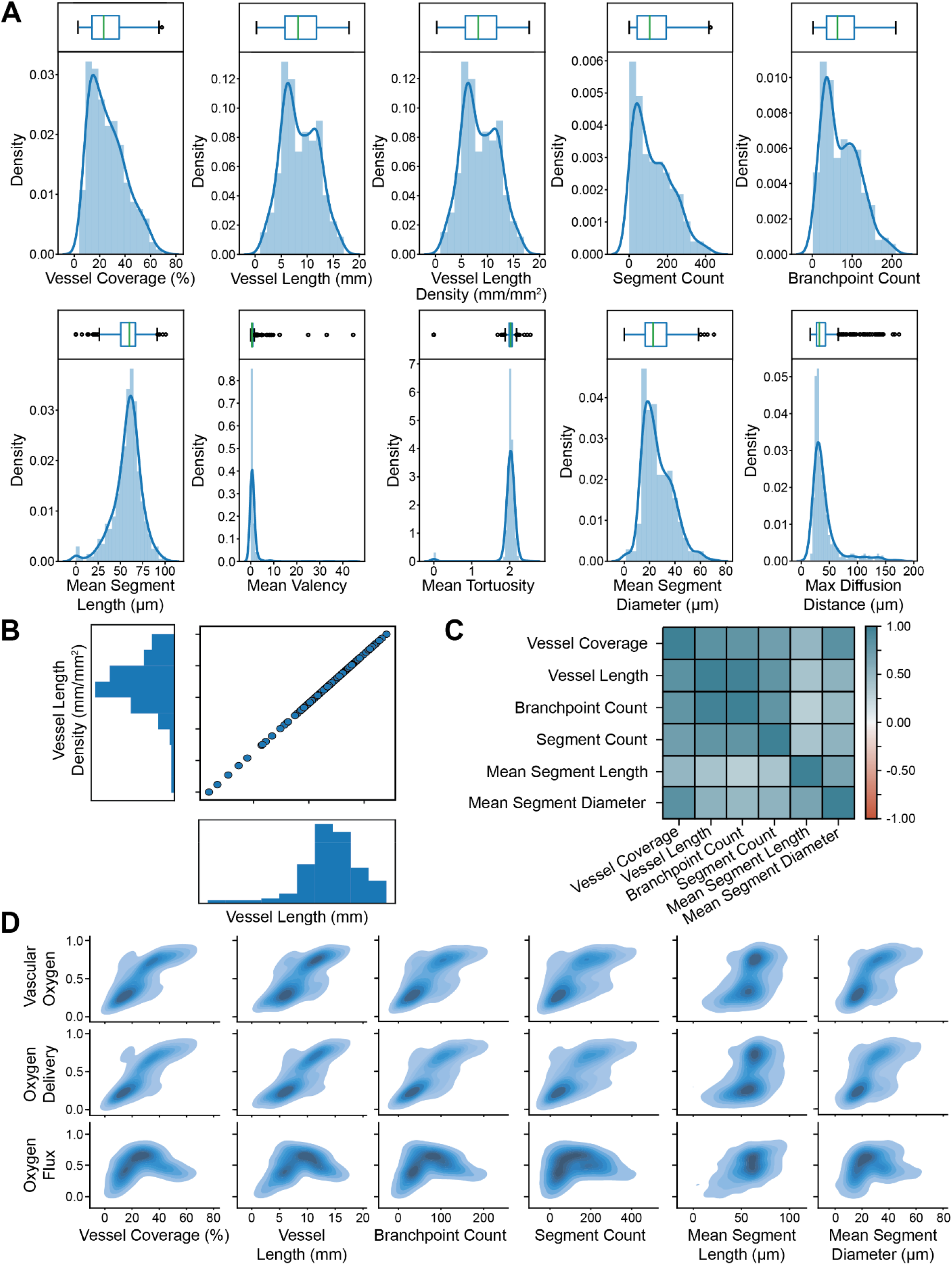
Exploratory data analysis aids in the generation of a suitable set of variables for downstream machine learning analyses. **A**) Histograms with overlaid kernel density estimation curves, topped with boxplots illustrating the mean, minimum and maximum, interquartile (Q1-Q3; 25-75%) range, and outliers. **B**) Distributions of vessel length density and vessel length. **C**) Heatmap of Pearson R test correlations among all morphological metrics. **D**) Scatter plots of morphological metrics against biological metrics for pattern recognition.

Input variables in machine learning models ought to be mutually exclusive and independent, or not multicollinear, in most machine learning algorithms. Traditionally, multicollinearity undermines the statistical significance of some independent variables, leading to a redundant multivariate equation, or may limit our ability to probe each of the independent variables’ impact on the predictive model^32,33^. Even though the variables were independent, we found through the Pearson R test that all these variables were highly correlated with each other, suggesting significant multicollinearity amongst some metrics (**Figure 2C**). We quantified the degree of multicollinearity by performing a Variance Inflation Factor (VIF) test (**Equation 1**)^34^, where a score above 5-10 indicates it being multicollinearly related to other variables in the dataset. Upon calculation, we found that all our input variables scored above this threshold, further confirming our concerns of multicollinearity (**Table S4**). Therefore, while independent but multicollinear variables can be used in some machine learning algorithms, it limits the reliability of some models.

To conclude our EDA, we generated scatter plots of each of our biological metrics against our morphological metrics for pattern recognition in our data (**Figure 2D**). We found that the features plotted against NVP and NOD showed density maps that may be fitted with simple geometries such as a linear or logarithmic curve, whereas five of the six same morphological metrics plotted against NOF resulted in parabolic shapes. Further, as NOF relies on the available oxygen from the vasculature (i.e. NVP) and quantifies an oxygen delivery rate as opposed to a mass of oxygen delivered (such as in NOD), we decided that a regression analysis might also benefit from only predicting NVP or NOD.

### Dimensional reduction of metrics: Principal component analysis and factor analysis

Since the metrics are multicollinear, we next set out to examine feature engineering to overcome multicollinearity. Dimensional reduction is a common method of engineering new features, and such an exercise is beneficial for applications such as multiple linear regression with interacting variables. Principal component analysis (PCA) is a common dimensional reduction technique that generates a new set of linearly combined variables representing decreasing amounts of variance among the entire dataset. For visualization, we classified each sample by labeling them as “Great”, “Good”, “OK”, and “Bad”, determined by splitting normalized biological values equally into four parts. After PCA, we saw a separation between “Great” and “Bad” samples in a scatter plot of Principal Component 1 (PC1) and Principal Component 2 (PC2) (**Figure 3A**). The two moderate groups localized in the middle with some overlap. We chose a 2-dimensional plot with only PC1 and PC2 because they accounted for a combined 88.45% of the total variance of the dataset, and the two had eigenvalues of 4.30 and 1.02, respectively (**Figure 3B-C**). These are traditional norms selecting for principal components (PCs), and we thus found it suitable to explain the entire dataset with only two PCs, or two variables^35^. Having two PCs suggests that a regression problem might be defined with two independent variables formed via linear combinations of the original morphological data. To support this, we generated a Pearson R test with the two PCs and the original morphological variables, finding that vessel coverage, vessel length, branchpoint count, and segment count loaded above a threshold of |0.4|^36^ on PC1, and mean segment length and diameter loaded on PC2 above the same threshold (**Figure 3D, Table S5**).

**Figure 3.**
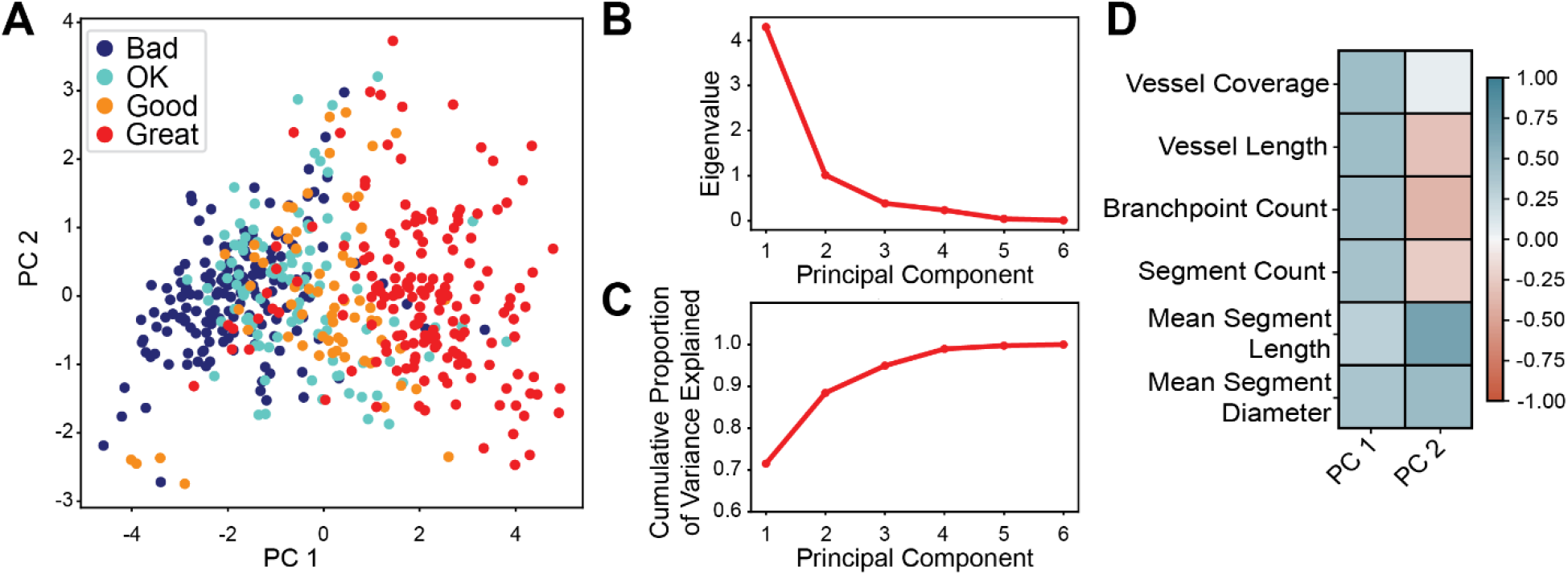
Dimensional reduction of variables with principal component analysis (PCA). PCA is used to determine relationships between all variables and linearly combined variables (PC1 and PC2) encompassing large proportions of variance within the entire dataset. **A)** The scatter plot between PC1 and PC2 with classification of data based on great, good, ok, or bad criteria. Scree plots showing the **B**) eigenvalue and **C)** cumulative proportion of variance, vs principal components. **D)** Correlation heatmap between all morphological variables and the PCs.

Exploratory factor analysis (EFA) is another technique used to determine potential relationships between the original, measurable variables (manifest variables) from our database that may result in new underlying variables (latent variables) that cannot be measured directly. Similar to PCA, exploratory FA may result in new variables that can dimensionally reduce the size of the dataset. To determine the adequacy of this analysis, we ran a Bartlett Sphericity Test, which tests the null hypothesis that the correlation matrix of the manifest variables is an identity matrix, which if not rejected, would mean that the variables are unrelated and unideal for factor analysis^37^. We observed a p-value < 0.001 indicating that the dataset is suitable for factor analysis. Additionally, we ran a Kaiser-Meyer-Olkin (KMO) Test, another adequacy benchmark measuring how the factors explain each other^37^. A value less than 0.5 indicates FA is unideal, and a value close to 1.0 indicates FA is ideal. We observed a KMO value of 0.71, again indicating FA could be applied in our study.

After FA, we observed that the first two output Factors contained eigenvalues above 1 (**Figure 4A**), a criterion determined due to this value representing the variance of at least one manifest variable^37^. Within these two factors, we found that all manifest variables loaded to one of the latent variables (Factor 1), while branchpoint count, mean segment length, and mean segment diameter loaded on the second (Factor 2) (**Figure 4B, Table S5**). These observations suggest that if new interacting factors were assembled to replace the original six morphological variables, the regression equation may still have redundancies due to some morphological values contributing to multiple Factors.

**Figure 4.**
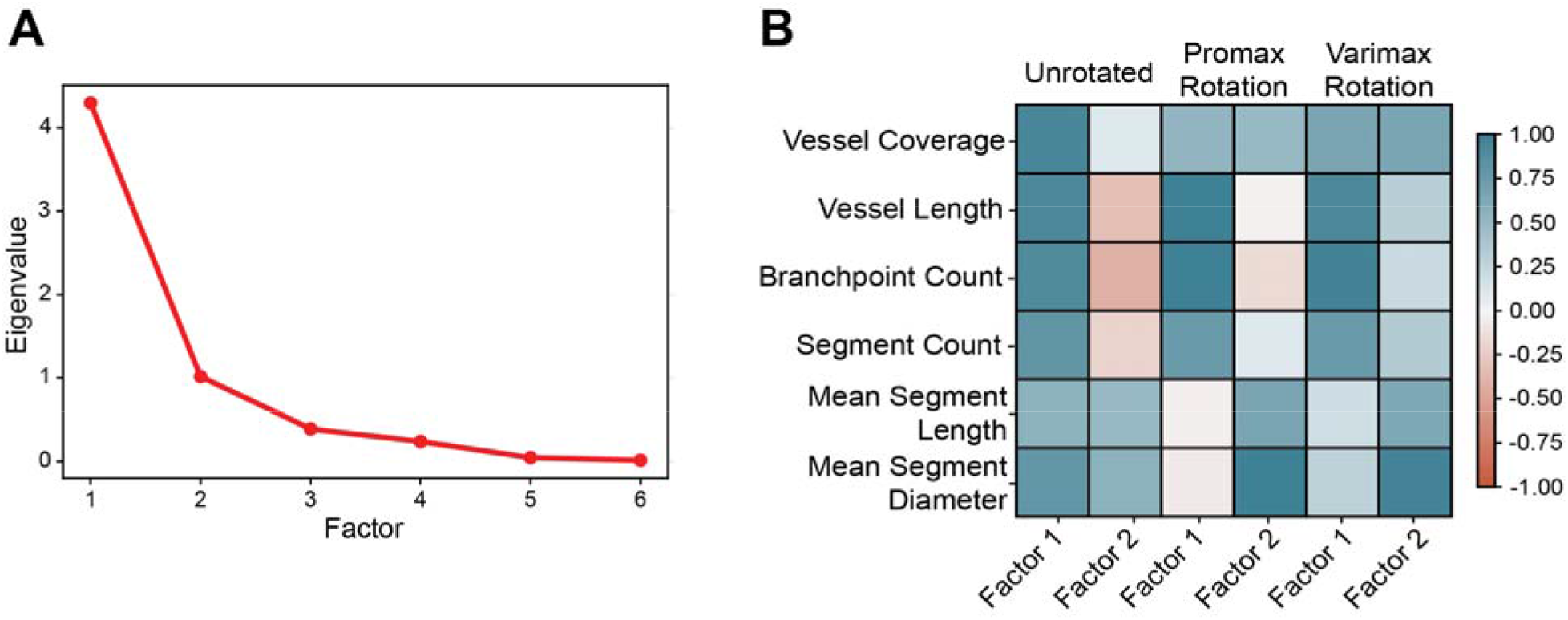
Dimensional reduction of variables with factor analysis (FA). **A**) Scree plot of eigenvalues vs factors, and **B)** Unrotated, Promax Rotated, and Varimax Rotated Factor Analyses segregate the morphological inputs based on their correlation values (i.e. loadings on each Factor).

Factor rotations are a common technique to decrease the complexity and increase the interpretability of the FA^37^. The rotations can be categorized as orthogonal or oblique. Orthogonal rotations may increase the simplicity of the FA model because the output loadings represent correlations between the new factors and the observed features, similar to our PCA results. Upon a Varimax (orthogonal) rotation, we found that vessel coverage correlated highly with both Factors, and the remaining 5 manifest variables were distributed to one or the other of Factor 1 and 2 (**Figure 4B, Table S5**). In an oblique rotation, the loadings represent regression coefficients. We used a Promax (oblique) rotation, which resulted in different loading values but the same assignment of morphological variables as the Varimax rotation (**Figure 4B, Table S5**). Therefore, these analyses suggested that it might be possible to explain the entire dataset with three variables: vessel coverage, Factor 1 (the combination of vessel length, branchpoint count, and segment count), and Factor 2 (the combination of mean segment length and mean segment diameter). This is slightly different from the PCA results because vessel coverage only significantly loaded on PC1, whereas after FA, vessel coverage significantly loaded on both factors and thus should remain as its own variable in a regression problem with interacting variables.

Finally, when we again analyzed the multicollinearity of the dataset created using vessel coverage, Factor 1, and Factor 2, we found a significant reduction in VIF values, although they were still above the threshold of mutual independence (**Table 6S**). Taken together, our analysis shows that a combination of vessel length, branchpoint count, and segment count, and a combination of mean segment length and mean segment diameter, represents higher independence and lower multicollinearity than these individual variables as inputs in a higher dimensional model.

### Machine learning applied to vascular networks: Multiple linear regression

After our evaluation of the input metrics, multiple linear regression (MLR) emerged as an obvious first candidate for quantitating the vascular networks using machine learning. Also, based on the positively linear shapes of the scatter plots between each of the independent and dependent variables (**Figure 2D**), we determined that a linear regression rather than logistical, logarithmic, or exponential, is more appropriate. We split the dataset randomly into two sub-groups: a training set (80% of the entire dataset) to train our machine learned model, and a test set (20% of the entire dataset) to evaluate the accuracy of the model. We decided that standard accuracy metrics such as the R^2^ value, mean absolute error (MAE), mean squared error (MSE), and root mean squared error (RMSE) would be adequate for evaluating the performance of our regression models.

For each dependent variable (NVP or NOD), we generated three MLR equations: 1) an equation with all six morphological values; 2) the same equation after eliminating insignificantly contributing morphological values via recursive feature elimination with cross validation (RFCEV), and; 3) an equation using vessel coverage, Factor 1, and Factor 2 from the feature engineering experiment. As the equations become more simplified, we expected the elimination of insignificantly contributing variables would increase the accuracy of the regression equation. Further, we were interested in examining whether the engineered, less multicollinearly related features might result in more accurate predictions of each of NVP and NOD. Finally, before generating the regression equations, all morphological values were scaled to dimensionless units between 0-1 to aid in later analyses determining feature importance via regression coefficient sign and magnitude.

The two MLR equations resulted in models with good predictive performance shown qualitatively in scatter plots of the predicted vs actual values after testing (**Figure 5A-C**) and quantitatively with the traditional accuracy metrics (**Table S7**). An MLR equation using all input features generated models predicting the NVP (**Figure 5A-i**) and NOD (**Figure 5A-ii**) with moderate accuracy, where the models had an R^2^ value of 0.64 and 0.74 and a MAE of 0.14 and 0.09 for NVP and NOD, respectively (**Table S7**).

**Figure 5.**
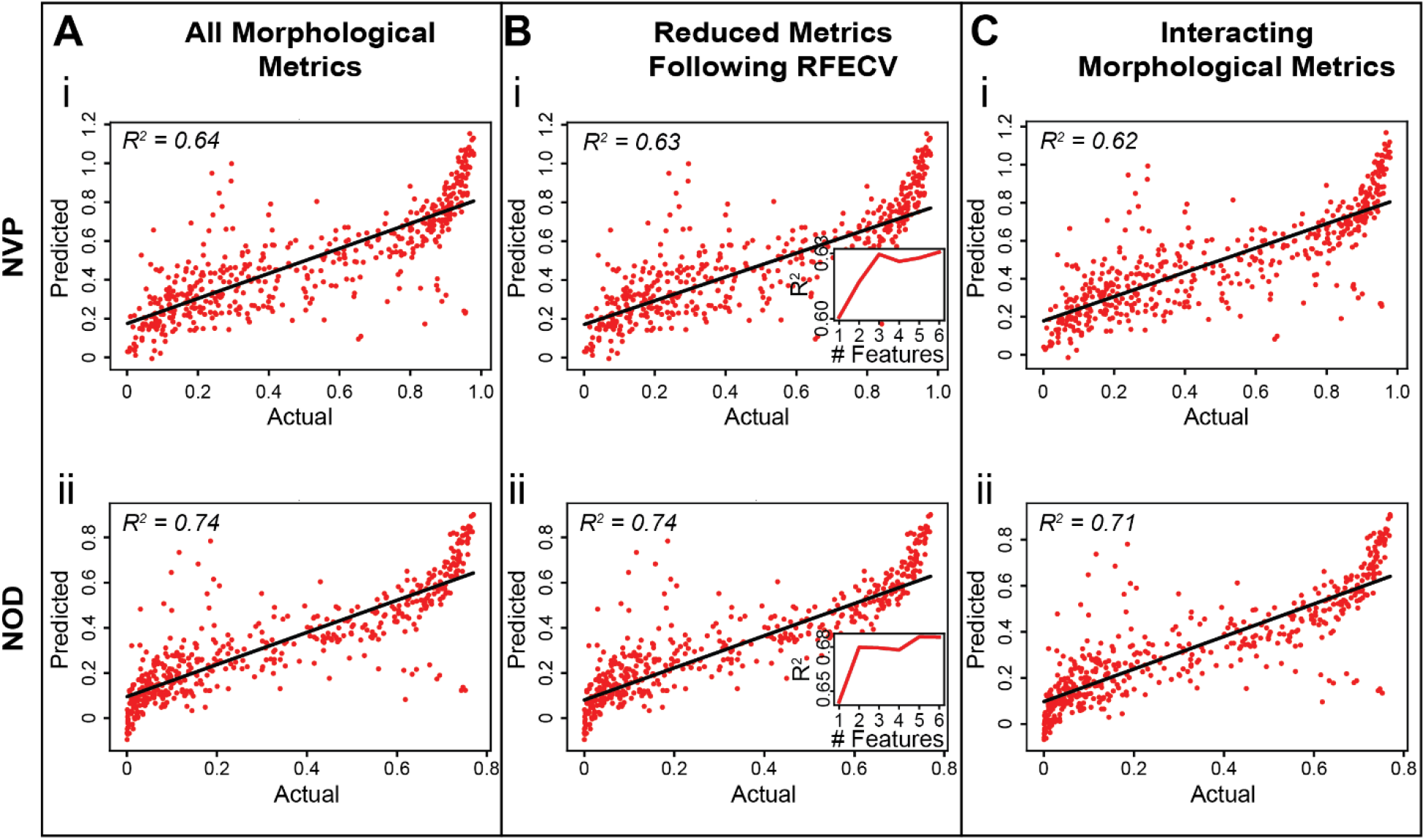
Machine learned multiple linear regression shows moderate success in predicting biological metrics. Analyses include **A**) regression using all 6 morphological metrics; **B**) regression following recursive feature elimination with-cross validation (RFCEV), and; **C**) regression analysis consisting of combinatorial variables from exploratory FA (vessel coverage, Factor 1, and Factor 2). Regression analyses using A-C-i) Normalized Vascular Potential as the dependent variable, and; A-C-ii) Normalized Oxygen Delivery as the dependent variable.

Interestingly, RFECV eliminated a single morphological variable in the NVP equation (**Figure 5B-i**), but no variables in the NOD equation (**Figure 5B-ii**), highlighting that our feature selection experiment yielded a good, non-redundant set of inputs. Eliminating a single variable from the NVP equation resulted in a slight and insignificant decrease in accuracy. Also, models using interacting, combinational factors performed similar to the models using non-interacting variables (**Table S7, Figure 5Ci-ii**), highlighting that dimensional reduction reduces prediction time via a lower dimensional model, but similarly performing models using raw morphological data may still be used without this intermediate step.

Simple observation of the regression coefficients is a common method of determining which of the input metrics influences the prediction. However, the multicollinearity of our data makes this analysis limited due to the mutual dependence amongst the variables. Nevertheless, we found that the regression coefficients for vessel coverage outweighed all others in every case by 2-3-fold (**Table S8**), but the next most impacting metric differs for each biological variable. For example, vessel length gives a slightly negative coefficient for NVP, suggesting higher vessel lengths can decrease the available oxygen to diffuse into the hydrogel region. This observation is not true for NOD, however, as vessel length was the least contributing variable with a regression coefficient of 0.02, causing its elimination during RFECV. Finally, we observed that the morphological data explains the highest amount of variance among the NOD data (71-74%), shown by the R^2^ scores (**Figure 5, Table S7**), confirming our original hypothesis that vascular morphology may describe vascular function. Thus, this model is a quick and simple method of predicting AngioMT values from morphological data with ±15% error, and suggests vessel coverage to be most predictive of a highly functional vascular network.

### Machine learning applied to vascular networks: Random forest regression

Decision tree based-regression is a machine learning technique that is not limited by multicollinearity, which our dataset exhibits. Hence, we first established a decision tree regressor, despite their known susceptibility to overfitting of the training dataset, which limits their power with unknown data. Expectedly, the decision tree regression model showed poor accuracy when undergoing testing (**Table S9**). However, the residuals show a normalized distribution centered at 0 for each biological metric (**Figure 6A-i**). Interestingly, a scatter plot of the expected vs. predicted oxygen delivered shows that higher values give higher errors, but lower values tend to be predicted accurately (**Figure 6A-ii**). Further, vessel coverage is the most important metric (**Figure 6A-iii**), proven via a computation of the mean decrease in accuracy of the tree as each metric is randomly permutated, a common technique for feature importance identification. Similar to MLR, the other morphological metrics contribute to each prediction at least 2-fold less.

**Figure 6.**
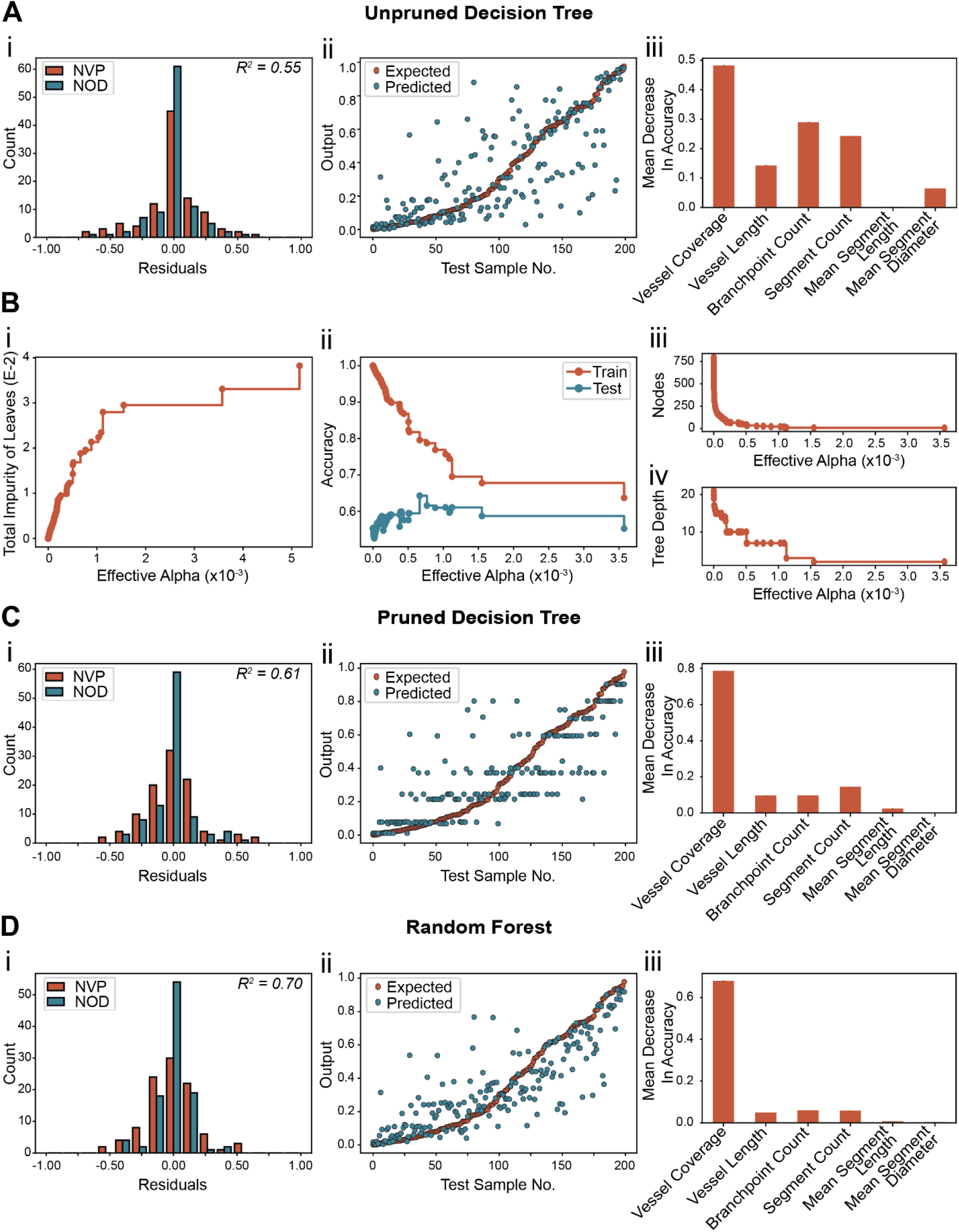
Tree-based regression methods demonstrate comparable accuracies but overcome problems of multicollinearity. **A**) Unpruned decision tree testing results. i) Residuals histogram. ii) Expected vs. predicted scatter plots. iii) Random permutation of input variables to measure the chief governor of the model. **B**) Results of testing to generalize the overfit, unpruned tree. i) Tuning effective alpha to increase leaf impurity. ii) Training and testing data performance as effective alpha is increased. iii) Node number as effective alpha is increased. iv) Tree depth as effective alpha increases. **C**) Accuracy results after pruning the decision tree i) Residuals histogram. ii) Expected vs. predicted scatter plot. iii) Random permutation of the input variables to determine the most important input. **D**) Random forest regressor testing results. i) Residuals histogram. ii) Expected vs. predicted scatter plots. iii) Random permutation of inputs to determine greatest contributor to the model.

A common technique to improve the accuracy of a decision tree regressor is pruning. Instead of arbitrarily limiting the depth of the tree or the number of leaves, we methodically pruned using a cost-complexity analysis^38^. Here, we used a tree score employing a penalty score (Effective Alpha), that characterizes the extent of pruning, which generalizes the tree but also increases the impurity of the leaves (**Figure 6B-i**), or decreases the accuracy of the model during training (**Figure 6B-ii**). Further, as we prune more of the tree by increasing the effective alpha, we decrease the number of nodes (**Figure 6B-iii**) and the depth of the tree (**Figure 6B-iv**), thus making a more generalized tree that might predict the testing data better than an unpruned tree.

Indeed, upon evaluating the best performing pruned decision tree regressor with the unseen testing data, we found that the accuracy metrics improve (**Table S9**) and the histogram of residuals remained normally distributed around 0 (**Figure 6C-i**). A scatter plot of the expected vs. predicted values showed that the predicted data arranged in tiers, a common outcome of a pruned decision tree regressor (**Figure 6C-ii**). Strikingly, random permutation of vessel coverage tended to decrease the accuracy almost 80%, confirming that this metric influences the regression model (**Figure 6C-iii**).

Lastly, we employed random forest regression, an algorithm that employs a large ensemble of different, uncorrelated regression trees operating in committee that outperforms any single tree. Following training and subsequent testing, we found that a random forest performed significantly better than the previous tree architectures, with an R^2^ value of 0.70 (**Table S9, Figure 6D**). We again generated a histogram of the residuals, finding that the random forest model had the smallest range distributed normally around an error of 0 (**Figure 6D-i**). The expected vs. predicted scatter plot also showed that predicted values followed the expected trend to a closer extent than other tree-based regression models, yet some outliers still emerge (**Figure 6D-ii**). Finally, an analysis of the mean decreases in accuracy following random perturbation of each morphological metric showed that vessel coverage was again the most important metric for this model (**Figure 6D-iii**). Therefore, random forest regression is a predictor of AngioMT data using morphological data as input, even when multicollinear data is presented. Further, random forest regression outperforms simpler tree-based regressors, and may thus be adopted for approximating AngioMT quantifications of the vMPS without dedicating high computational resources to their image processing pipeline.

## 4. Discussion

This exercise in exploratory data analysis, dimensional reduction, and superficial machine learning provides a framework for designing vMPS with a biological motivation. Our key findings include a reduction of morphological variable inputs into the model, two new linearly combined variables encompassing the morphology with lower multicollinearity, and evidence that machine learning algorithms may be able to assist in predicting the biological function of capillary networks cultivated in vMPSs. Importantly, random forest regression emerged as an optimal algorithm for predicting oxygenation given the nature of our morphological data. This model is now available for future vMPS engineers to optimize their networks without having to run their own numerical analyses to measure biological metrics of oxygen transport.

Our analysis also revealed that among all singular metrics used, vessel coverage is a morphological measure of vascular networks that is statistically most relevant. However, exclusively relying on the vessel coverage could lead to improper outcomes from vMPS. For example, Kim, *et al*., found that some vMPS exhibited characteristics of hyperplasia and subsequently engineered thinner networks with lower vessel coverages via pericyte co-culture^39^. They found that pericytes reduced the vessel diameter and increased the number of junctions and branches, and even decreased the vessel permeability. Thus, the other five morphological metrics included in our own analyses might still play an important role in the biologically motivated engineering of vascular networks in MPS. Further, for the remaining five morphological metrics, RFECV after linear regression found that the other five morphological metrics generally increased the accuracy of the predictions, and random feature perturbation in tree models showed some impact on the prediction accuracy. Therefore, it is still important to include all six metrics when predicting oxygen delivering potential.

This exercise also represents a need for the MPS and vMPS fields to begin adopting advanced quantification methods. We see that with simple programming, we can extract a large amount of data that relates a computationally inexpensive set of morphological metrics to a more computationally taxing numerical analysis of the potential oxygen transport and delivery through the same samples. As more complex assays are developed, especially towards the use of MPSs in medical regulation, pharmaceutics, and tissue engineering, more complex data analysis techniques may be adopted. Further, as more data is generated, machine and deep learning methods may be employed over manual data analysis^40,41^. Machine learning models can also be shared between labs, offering a way of standardizing MPS design procedures, such as the establishment of vMPS with tissue-specific organoids. As MPSs continue to increase in complexity, similar exercises in machine learning can be undertaken following the examples we completed herein.

## 5. Acknowledgements

This material is supported by an American Heart Association Predoctoral Fellowship under Grant No. 906239, a National Science Foundation Graduate Research Fellowship under Grant No. 1650114, and a Texas A&M Biomedical Engineering Department National Excellence Fellowship to J.J.T.; and by the NHLBI of NIH under Award Number R01HL157790, NSF CAREER Award number 1944322, and Texas A&M University President’s Excellence in Research Award (T32/ X-Grant) to A.J.

