## Supplemental Information for "Evaluation of the morphological and biological functions of vascularized microphysiological systems with supervised machine learning"

**Supplementary Methods**

*Microfluidic Device Fabrication*

Microfluidic devices were manufactured according to traditional photo- and soft-lithography practices. First, photomasks containing multiple iterations of the microfluidic device were designed using Solidworks 2019 (Dassault Systems). Individual devices consisted of 5 parallel channels separated by rows of hexagonal microposts. Channels serving as hydrogel compartments had a width of 1.5 mm while fluidic channels were 0.5 mm wide. All channels were 8 mm long and terminated by 1 mm circular ports for hydrogel channels and 5 mm circular ports for fluidic channels. Finalized photomask designs were sent out for printing (CAD-Art Services; Bandon, Oregon, USA).

SU-8 2100 photoresist (Kayaku Advanced Materials; Westborough, MA 01481) was spin coated onto 6 in silicon wafers (University Wafer; South Boston, MA 02127, USA) to a height of 125 µm and baked at 95 °C for 60 min. The process was repeated resulting in a SU-8 height of 250 µm. SU-8-coated wafers were placed in a UV-lamp box (BlackHoleLab; Paris 75011, France) and covered by the mylar photomask before exposure to UV. UV-treated wafers were baked at 95 °C for 20 min and developed in a shaking bath of SU-8 developer (Kayaku Advanced Materials) for 20 min. Developed wafers were then washed with more developer followed by isopropanol and subsequent air drying via compressed air gun. Finished wafers were then silanized with trichloro(1H,1H,2H,2H-perfluorooctyl)silane (PFCOTS) (448931-10G, Sigma Aldrich) overnight.

Polydimethylsiloxane (PDMS) (Dow Corning) was prepared by mixing the polymer base with the curing agent at a 10:1 ratio using an industrial mixer. Combined pre-polymer was then poured onto the master mold wafer and degassed for 15 min before baking at 70 °C for 2 hr. Cured PDMS was peeled from the master mold and port holes were punched using 1 and 5 mm biopsy punches. Packaging tape was used to remove debris from around the port holes before subjecting the cut PDMS slabs to a short burst of compressed air. Devices along with glass microscope slides were placed facing up in a 100 Watts plasma cleaner (Thierry Zepto, Diener Electronics; Ebhausen 72224, Germany). Following treatment, devices were compressed onto the slides to facilitate irreversible bonding and left at 70 °C overnight to render the PDMS hydrophobic. Before cell culture, devices were left under UV for 30 min for sterilization.

*Numerical Analysis*

Sectioned, binary images from each sample’s vascularization region (image size: 1000 x 1000) were imported in MATLAB and converted to triangular meshes using im2mesh, an image to mesh converter developed by Jiexian Ma. Meshes were generated for the non-vascularized hydrogel regions (tissue phase) as well as for vascularized networks (vessel phase). Once a composite triangular mesh was obtained, we developed AngioMT, a Galerkin finite element method-based mass transport solver in MATLAB for studying the two-dimensional mass transport of oxygen through the vascularized networks. AngioMT first detected edge nodes between the vessel and tissue phases, which were later used to apply boundary conditions. Oxygen inlets were defined at the left and right extremes of each image i.e., at x=0 and x=1000, and an oxygen concentration value equivalent to the oxygen concentration inside a humidified 5% CO_2_ incubator were applied as inlet boundary conditions. AngioMT then applied an oxygen permeability flux at the edges signifying the transport of oxygen from vessels to the surrounding tissues (hydrogels in this case). Once these boundary conditions were applied, AngioMT solved the steady-state two-dimensional transport of oxygen within the vessel phase. Once the nodal oxygen values were obtained, AngioMT then calculated the nodal oxygen concentration values within the tissue phase assuming a uniform oxygen consumption via a chemical reaction with the nodal oxygen values at the edges (obtained in the previous step) as the inlet boundary condition. After obtaining the nodal oxygen concentration values in both phases, AngioMT calculated the area averaged oxygen concentration within the vessel and tissue phases and reported them as Normalized Vascular Potential and Normalized Oxygen Delivery respectively, after normalization with respect to the oxygen concentration at inlets.

*Data Analysis and Machine Learning*

All data analyses were carried out using Python 3.8.8 via Anaconda3, and all code was written in JupyterLab 3.0.14 web browser. REAVER datasets were modified with an additional column for NOF, NVP, and NOD measurements corresponding to each sample and used as input for all analyses.

Exploratory data analysis was first conducted on the entire dataset consisting of all output variables from REAVER and the in-house generated NOF, NVP, and NOD values. Histograms were generated using pandas hist(), and correlation heatmaps were made using seaborn heatmap() with an argument for correlation values generated from pandas corr(). Scatter plots were generated using seaborn pairplot(). Following this, the following variables were excluded from all subsequent analyses: Vessel Length Density, Mean Valency, Mean Tortuosity, and Max Diffusion Distance.

VIF was calculated to quantify the degree of multicollinearity between the remaining morphological metrics using variance_inflation_factor() from statsmodels stats.outliers_influence().

For PCA, all data except NVP and NOD, which have a range of 0.0-1.0, was scaled using sklearn MinMaxScaler(). PCA() from sklearn.decomposition was used with the components argument set to 6, thus generating 6 principal components explaining a decreasing proportion of variance. For visualization’s sake, the new PCs were segregated based on their input NOD value: Great – 1.0-0.75; Good – 0.50-0.75; OK – 0.25-0.50; Bad – 0.0-0.25. Scatter plots of PC1 vs. PC2 labeled by their NOD classification were generated using ax.scatter(). Scree plots showing the proportion of variance explained by each PC were by calling PCA.explained_variance_ratio in a matplotlib plot(), and eigenvalues from which the number of PCs were chosen were determined by calling PCA.explained_variance. Loadings were obtained using PCA.comonents_. Heatmaps of PCs vs. metrics were generated as previously described.

Input data for FA was scaled similar to the PCA methods. Bartlett Sphericity and KMO tests were generated using calculate_bartlett_sphericity() and calculate_kmo() from factor_analyzer.factor_analyzer. Unrotated FA was first carried out using FactorAnalyzer from factor_analyzer with the number of factors set to 6. Loadings were obtained using FactorAnalyzer.loadings_, and eigenvalues were retrieved from FactorAnalyzer.get_eigenvalues(). Scree plots were generated similar to the PCA methods, from which we determined to use only the first two factors. For rotations, we re-ran FactorAnalyzer() with an input argument for rotations set to ‘varimax’ for the Varimax rotation and ‘promax’ for the Promax rotation.

Data for machine learning was split using scikit-learn train_test_split (), which separated the x and y data into a training set containing 80% of the data and a testing set containing the other 20%. LinearRegression() from sklearn.linear_model was used to fit the training dataset, and LinearRegression.predict() was used to generate the predicted biological values with the trained model. Accuracy metrics were determined using sklearn.metrics (r2_score(), mean_absolute_error(), and mean_squared_error()). Numpy polyfit() was used to generate the line of best fit for the expected vs. predicted plots, giving values for the slope and y-intercept. Scatter plots were generated using previously mentioned methods while stacking the line of best fit over the scatter plot. To further probe the performance of the multiple linear regression model, we used RFECV() from sklearn.feature_selection, which eliminates the least important feature before re-running the regression iteratively until the accuracy of the model decreases in accuracy.

Decision tree regression was carried out using DecisionTreeRegressor() from sklearn.tree. Accuracy metrics were determined similar to multiple linear regression methods. The pruned decision tree regressor was generated by determining the optimal effective alpha value via a cost-complexity analysis that aimed to mitigate the effects of overfitting and maximizing accuracy on the testing dataset, accomplished using cost_complexity_pruning_path() from DecisionTreeRegressor, with effective alpha values recalled from cost_complexity_pruning_path.ccp_alphas() and impurities from cost_complexity_pruning_path.impurities(). For the best performing pruned decision tree regressor, we chose an alpha value resulting in the smallest accuracy difference between the training and testing dataset. The pruned decision tree model was generated by using DecisionTreeRegressor() with an input argument for the optimal ccp_alpha

Random forest models were generated using RandomForestRegressor() from sklearn.ensemble. An input argument for n_estimators was set to 100, resulting in a random forest model consisting of 100 unique and uncorrelated decision tree regressors. The feature importances represented as decreases in accuracy following morphological metric perturbation were recalled from the model using permutation_importance() from sklearn.inspection.

All plots were exported as SVG files using pyplot.savefig from matplotlib and developed into publication-ready figures using Adobe Illustrator (Adobe, Inc.; San Jose, CA 95110, USA).

**Supplementary Tables**

**Table S1** – Morphological and functional metrics used to describe vMPSs in this study.

| *Variable* | *Units* | *Description* |
| --- | --- | --- |
| **Vessel Coverage** | **%** | Fraction of pixels belonging to blood vessels within an image |
| **Vessel Length** | **mm** | Centerline length of vessels within an image |
| **Vessel Length Density** | **mm^2^/mm** | Centerline length of vessels normalized by the image area |
| **Branchpoint Count** |  | Number of branchpoints calculated from the vessel centerline |
| **Segment Count** |  | Number of segments calculated from the vessel centerline, including between two branchpoints, from a branchpoint to a terminal point, and from a branchpoint to the edge of an image) |
| **Mean Segment Length** | **µm** | Average length of segments within an image |
| **Mean Tortuoisty** |  | Image averaged segment complexity score representing the distance between a segment start and end point |
| **Mean Valency** |  | Score representing the complexity of a vascular network, calculated as the ratio of the branchpoint and segment counts |
| **Mean Segment Diameter** | **µm** | Average diameter of segments within an image |
| **Max Diffusion Distance** | **µm** | Image averaged measurement of the average distance between a point in the tissue and its nearest vessel |
| **Normalized Oxygen Flux** |  | Average oxygen diffusion rate from vessels to tissues normalized by inlet oxygen concentration |
| **Normalized Vascular Potential** |  | Area average oxygen concentration of the vasculature normalized by inlet oxygen concentration |
| **Normalized Oxygen Delivery** |  | Area average oxygen concentration of the tissue normalized by inlet oxygen concentration |

**Table S2** – statistical descriptors of the entire dataset.

| *Metric* | *Mean* | *Standard Deviation* | *Minimum* | *25th Percentile* | *50th Percentile* | *75th Percentile* | *Maximum* |
| --- | --- | --- | --- | --- | --- | --- | --- |
| **Vessel Coverage** | 26.4148 | 13.908063 | 3.8354 | 14.825625 | 23.8877 | 35.978675 | 69.3088 |
| **Vessel Length** | 8.5499 | 3.385138 | 0.97152 | 5.928847 | 8.182056 | 11.35143 | 17.016 |
| **Vessel Length Density** | 8.5499 | 3.385138 | 0.97152 | 5.928847 | 8.182056 | 11.35143 | 17.016 |
| **Branchpoint Count** | 72.378 | 44.953852 | 2 | 36 | 63.5 | 105.25 | 209 |
| **Segment Count** | 126.684 | 97.512078 | 0 | 40.75 | 107.5 | 193.25 | 426 |
| **Mean Segment Length** | 57.8833 | 15.667722 | 0 | 50.288877 | 60.027036 | 66.788457 | 100.8 |
| **Mean Tortuosity** | 1.99995 | 0.266221 | 0 | 1.987365 | 2.027485 | 2.065635 | 2.549333 |
| **Mean Valency** | 1.11388 | 2.919014 | 0 | 0.433136 | 0.557536 | 0.871249 | 44.5 |
| **Mean Segment Diameter** | 25.562 | 11.771502 | 0 | 16.443155 | 22.689335 | 33.372863 | 70.698366 |
| **Max Diffusion Distance** | 41.4907 | 25.83976 | 15.882998 | 27.158085 | 32.090601 | 42.904593 | 173.685732 |

**Table S3** – Outlier calculations for each input variable

| *Metric* | *Number of Outliers* | *Proportion of Outliers* |
| --- | --- | --- |
| **Vessel Coverage** | 1 | 0.002 |
| **Vessel Length** | 0 | 0 |
| **Vessel Length Density** | 0 | 0 |
| **Branchpoint Count** | 0 | 0 |
| **Segment Count** | 1 | 0.002 |
| **Mean Segment Length** | 22 | 0.044 |
| **Mean Tortuosity** | 27 | 0.054 |
| **Mean Valency** | 57 | 0.114 |
| **Mean Segment Diameter** | 5 | 0.01 |
| **Max Diffusion Distance** | 51 | 0.102 |

**Table S4** – Variable inflation factor (VIF) calculations based on **Eqn. 1** as indicators of the degree of multicollinearity between each of the final list of morphological metrics chosen after exploratory data analysis.

| *Variable* | *VIF Value* |
| --- | --- |
| **Vessel Coverage** | 59.34 |
| **Vessel Length** | 146.87 |
| **Branchpoint Count** | 79.95 |
| **Segment Count** | 8.68 |
| **Mean Segment Length** | 36.14 |
| **Mean Segment Diameter** | 44.97 |

**Table S5** – PCA and EFA loadings for all variables above a threshold of |0.4|. PC loadings, FA loadings, and Varimax rotation loadings represent correlations between the Factors/PCs and each morphological metric. Promax rotation loadings represent regression coefficients for each variable.

| **Mean Segment Diameter** | **Mean Segment Length** | **Segment Count** | **Branchpoint Count** | **Vessel Length** | **Vessel Coverage** | *Morphological Metric* | |
| --- | --- | --- | --- | --- | --- | --- | --- |
| - | - | 0.41 | 0.43 | 0.44 | 0.40 | *PC 1* | *PC Loadings > \|0.4\|* |
| 0.47 | 0.68 | - | - | - | - | *PC 2* |  |
| 0.80 | 0.56 | 0.81 | 0.90 | 0.93 | 0.95 | *Factor 1* | *FA Loadings > \|0.4\|* |
| 0.55 | 0.49 | - | 0.41 | - | - | *Factor 2* |  |
| - | - | 0.75 | 0.97 | 0.93 | 0.67 | *Factor 1* | *Varimax Rotation Loadings > \|0.4\|* |
| 0.96 | 0.62 | - | - | - | 0.65 | *Factor 2* |  |
| - | - | 0.76 | 1.08 | 1.00 | 0.53 | *Factor 1* | *Promax Rotation Loadings > \|0.4\|* |
| 1.04 | 0.67 | - | - | - | 0.50 | *Factor 2* |  |

**Table S6** – VIF calculations of the dimensionally reduced dataset following PCA and EFA.

| *Variable* | *VIF Value* |
| --- | --- |
| **Vessel Coverage** | 15.11 |
| **Factor 1** | 11.50 |
| **Factor 2** | 7.64 |

**Table S7** – accuracy metrics of machine learned multiple linear regression models.

| *Accuracy Metric* | *Normalized Vascular*  *Potential* | | | *Normalized Oxygen*  *Delivery* | | |
| --- | --- | --- | --- | --- | --- | --- |
|  | *Raw* | *Post-RFECV* | *Interacting Factors* | *Raw* | *Post-RFECV* | *Interacting Factors* |
| **R^2^** | 0.64 | 0.63 | 0.62 | 0.74 | 0.74 | 0.71 |
| **MAE** | 0.14 | 0.14 | 0.15 | 0.09 | 0.09 | 0.10 |
| **MSE** | 0.03 | 0.03 | 0.04 | 0.01 | 0.01 | 0.02 |
| **RMSE** | 0.19 | 0.19 | 0.2 | 0.13 | 0.13 | 0.14 |

**Table S8** – RFECV rankings and regression coefficients of input morphological metrics for each multiple linear regression analysis using the biological metrics as the dependent variable.

| *Metric* | *Normalized Vascular Potential* | | | *Normalized Oxygen Delivery* | | |
| --- | --- | --- | --- | --- | --- | --- |
|  | *RFECV Ranking* | *Raw Regression Coefficients* | *Regression Coefficients Following RFECV* | *RFECV Ranking* | *Raw Regression Coefficients* | *Regression Coefficients Following RFECV* |
| **Vessel Coverage** | 1 | 0.87 | 0.87 | 1 | 0.74 | 0.74 |
| **Vessel Length** | 1 | -0.21 | -0.21 | 2 | 0.02 | - |
| **Branchpoint Count** | 1 | 0.42 | 0.42 | 1 | 0.25 | 0.27 |
| **Segment Count** | 1 | 0.17 | 0.17 | 1 | 0.14 | 0.14 |
| **Mean Segment Length** | 1 | 0.31 | 0.31 | 1 | 0.16 | 0.16 |
| **Mean Segment Diameter** | 1 | -0.13 | -0.13 | 1 | -0.16 | -0.16 |

**Table S9**– accuracy metrics of machine learned tree-based regression models.

| *Accuracy Metric* | *Decision Tree Regressor* | *Pruned Decision Tree Regressor* | *Random Forest Regressor* |
| --- | --- | --- | --- |
| **R^2^** | 0.55 | 0.61 | 0.70 |
| **MAE** | 0.12 | 0.13 | 0.11 |
| **MSE** | 0.04 | 0.30 | 0.02 |
| **RMSE** | 0.20 | 0.20 | 0.16 |
